# Clonal dissection of immunodominance and cross-reactivity of the CD4^+^ T cell response to SARS-CoV-2

**DOI:** 10.1101/2021.03.23.436642

**Authors:** Jun Siong Low, Daniela Vaqueirinho, Federico Mele, Mathilde Foglierini, Michela Perotti, David Jarrossay, Sandra Jovic, Tatiana Terrot, Alessandra Franzetti Pellanda, Maira Biggiogero, Christian Garzoni, Paolo Ferrari, Alessandro Ceschi, Antonio Lanzavecchia, Antonino Cassotta, Federica Sallusto

## Abstract

The identification of CD4^+^ T cell epitopes is essential for the design of effective vaccines capable of inducing neutralizing antibodies and long-term immunity. Here we demonstrate in COVID-19 patients a robust CD4^+^ T cell response to naturally processed SARS-CoV-2 Spike and Nucleoprotein, including effector, helper and memory T cells. By characterizing 2,943 Spike-reactive T cell clones, we found that 34% of the clones and 93% of the patients recognized a conserved immunodominant region encompassing residues S346-365 in the RBD and comprising three nested HLA-DR and HLA-DP restricted epitopes. By using pre- and post-COVID-19 samples and Spike proteins from alpha and beta coronaviruses, we provide in vivo evidence of cross-reactive T cell responses targeting multiple sites in the SARS-CoV-2 Spike protein. The possibility of leveraging immunodominant and cross-reactive T helper epitopes is instrumental for vaccination strategies that can be rapidly adapted to counteract emerging SARS-CoV-2 variants.

---

The prompt identification of T cell epitopes in disease causing organisms is a considerable challenge in view of the polymorphism of HLA class II molecules and the variability of rapidly mutating pathogens. Bioinformatic analysis is widely used to predict peptide binding to HLA molecules and to identify candidate targets of the T cell response (*1*). In the context of the COVID-19 pandemic, this approach has been used to predict T cell epitopes in SARS-CoV-2 proteins (*2, 3*) and to produce peptide pools that have been used to stimulate PBMCs in order to identify and enumerate antigen-specific T cells. These studies revealed a robust CD4^+^ and CD8^+^ T cell response against SARS-CoV-2 Spike, Membrane, Nucleoprotein and non-structural proteins in recovered patients (*2-6*) and a level of cross-reactivity with other circulating human coronaviruses in pre-pandemic samples (*7-9*).

A limitation of bioinformatic-based predictions is the difficulty to identify immunodominant epitopes, since immunodominance depends on multiple factors such as the constraints related to processing of whole proteins, the available T cell repertoire, HLA alleles and preexisting cross-reactive immunity (*10-12*). To identify naturally processed immunodominant CD4^+^ T cell epitopes we took an unbiased approach of stimulating memory CD4^+^ T cells with protein-pulsed professional antigen presenting cells (APCs), followed by the isolation of T cell clones that were used to precisely map the epitope recognized (*13*). In addition, TCR Vβ sequencing was used to define diversity and frequency of SARS-CoV-2-specific clonotypes within memory subsets and to estimate the extent of cross-reactivity with other coronavirus Spike proteins.

To analyze the CD4^+^ T cell response elicited by SARS-CoV-2, we collected blood samples from patients who had recovered from mild to severe COVID-19 disease 43-164 days post symptoms onset or positive PCR test (Table S1). From PBMCs of a first cohort of 14 patients, we isolated total CD4^+^ memory T cells or CD4^+^ T central memory (Tcm), T effector memory (Tem) and T circulating follicular helper (cTfh) cells (Fig. S1A). The cells were labelled with CFSE and stimulated with autologous monocytes in the presence of recombinant SARS-CoV-2 Spike or Nucleoprotein. In all patients we observed a strong proliferative response towards both antigens, as measured on day 6 by CFSE dilution, upregulation of ICOS and CD25 and production of IFN-γ, the latter being primarily detected in the Tcm and Tem subsets (**Fig. 1A, B** and **Fig. S1B, C**). The proliferative response was overall comparable in the three memory subsets, consistent with priming of both inflammatory Th1, helper T and memory T cells. In contrast, in unexposed individuals, the CD4^+^ memory T cell response to SARS-CoV-2 proteins was low or undetectable (**Fig. 1B** and **Fig. S1C**), consistent with the presence of a few cross-reactive T cells primed by seasonal coronaviruses (*4, 5, 9*).

**Figure 1.**
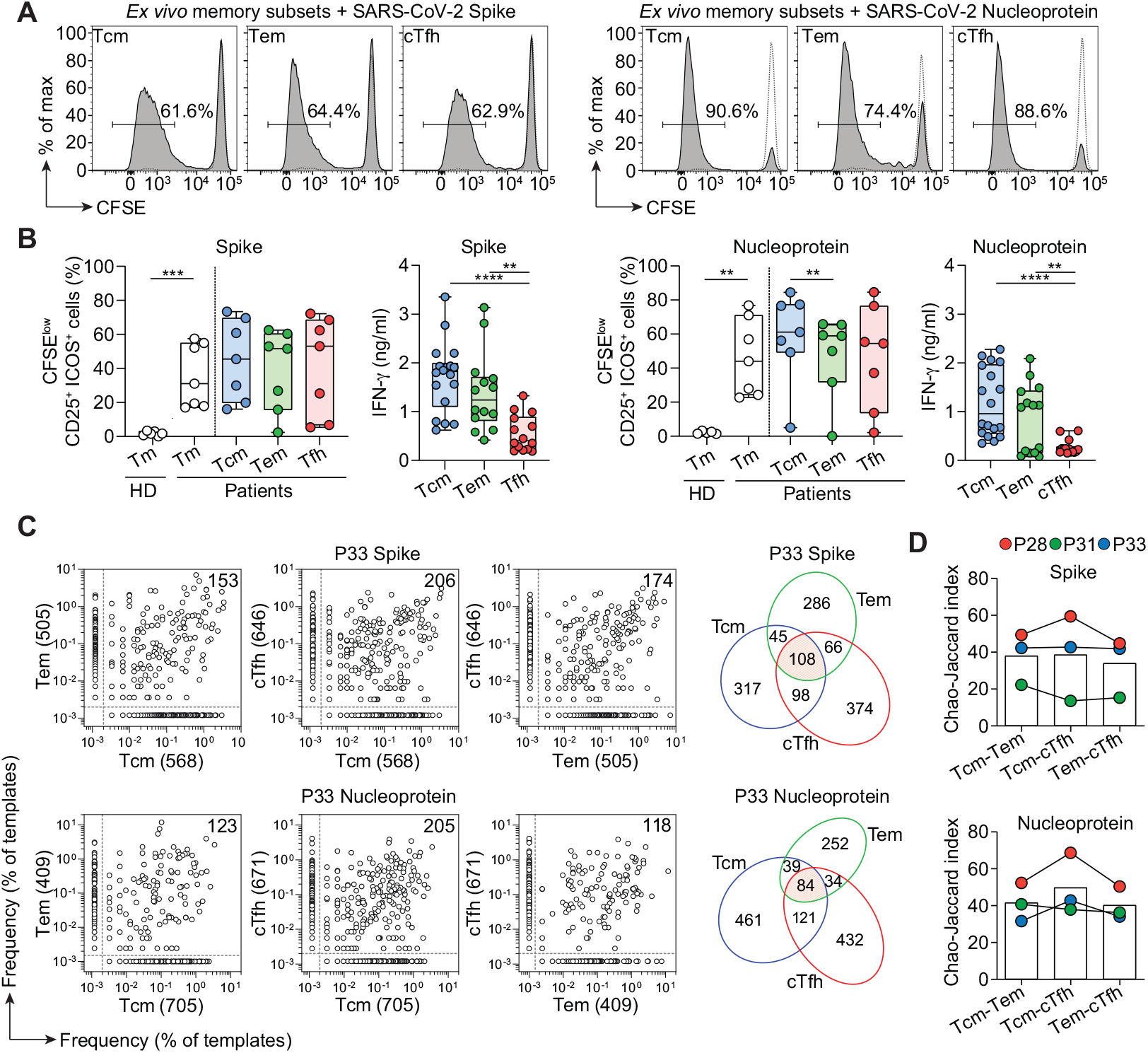
Robust T cell response to SARS-CoV2 Spike and Nucleoprotein in CD4^+^ memory T cell subsets. Total CD4^+^ memory T cells from 7 COVID-19 recovered patients and 6 unexposed (pre-COVID-19) healthy donors (HD) or CD4^+^ Tcm, Tem and cTfh cells from 7 COVID-19 recovered patients were labelled with CFSE and cultured with autologous monocytes in the presence or absence of recombinant SARS-CoV-2 Spike or Nucleoprotein. (**A**) CFSE profiles on day 7 and percentage of CFSE^lo^ proliferating Tcm, Tem and cTfh cells in a representative patient. Negative controls of T cells cultured with monocytes in the absence of antigen are reported as dashed lines. (**B**) Individual values, median and quartile values of percentage of CFSE^lo^CD25^+^ICOS^+^ cells in total CD4^+^ memory and CD4^+^ Tcm, Tem and cTfh memory subsets in the patients and healthy donors analyzed. Shown are also IFN-γ concentrations in the day 7 culture supernatants of SARS-CoV-2 Spike- or Nucleoprotein-stimulated memory T cell subsets. ****, *P* value < 0.0001 ***, *P* value < 0.001 **, *P* value < 0.01 as determined by two-tailed unpaired *t* test (total CD4^+^ memory and IFN-γ) or by two-tailed paired *t* test (CD4^+^ Tcm, Tem and cTfh). (**C**) Pairwise comparison of TCR Vβ clonotype frequency distribution in samples of T cells isolated from Spike-stimulated Tcm, Tem or cTfh subsets (initial input cells, 5×10^5^/subset) from patient P33. Frequencies are shown as percentage of productive templates. Total number of clonotypes is indicated in the *x* and *y* axis. Values in the upper right corner represent the number of clonotypes shared between two samples. The Venn diagrams show the number of unique and shared Spike- or Nucleoprotein-reactive clonotypes between Tcm, Tem and cTfh subsets of patient P33. The red area represents clonotypes identified in all 3 subsets. (**D**) Bar histograms showing the Chao-Jaccard overlap between pairs of TCR Vβ repertoires in patients P28, P31 and P33.

The clonal composition of SARS-CoV-2-reactive memory T cell lines and the relationship between different memory subsets was studied in three patients (P28, P31 and P33) by TCR Vβ sequencing of Spike-stimulated and Nucleoprotein-stimulated CFSE^lo^ T cell lines generated from each of the memory subsets. The Tcm, Tem and cTfh cell lines generated from the three patients comprised on average 908, 480 and 697 Spike-reactive clonotypes and 1,452, 623 and 908 Nucleoprotein-reactive clonotypes, respectively (**Fig. 1C** and **Fig. S2**). Interestingly, several of the most expanded clonotypes were shared between two and even among all three subsets, as shown by a high similarity index (**Fig. 1C, D**). Collectively, these findings reveal a robust polyclonal CD4^+^ T cell response to SARS-CoV-2 antigens. The polyfunctional response and the sharing of clonotypes among Tcm, Tem and cTfh cells is consistent with previous studies on the lineage relationship of CD4^+^ T cell subsets and with the intraclonal diversification of antigen primed CD4^+^ T cells (*14, 15*).

In view of the interest in vaccine design, we analyzed in depth the CD4^+^ T cell response to the Spike protein and in particular to the RBD, which is the main target of neutralizing antibodies (*16, 17*). CD4^+^ memory T cells from a larger cohort of 34 COVID-19 recovered patients (**Table S1**) were stimulated with Spike protein-pulsed APCs and proliferating T cells were isolated by cell sorting and cloned by limiting dilution. Using this approach, we obtained 2,943 T cell clones and mapped their specificity using 3 pools of 15-mer peptides overlapping of 10 spanning the S1-325 and S536-685 sequences (pool S1_ΔRBD_, comprising 91 peptides), the S316-545 sequence (pool RBD, comprising 44 peptides) and the S676-1273 sequence (pool S2, comprising 118 peptides) (**Fig. 2A, B**). Most donors analyzed (32 out of 34) had T cell clones specific for the RBD region (**Fig. 2B**). Further experiments using recombinant RBD to stimulate CD4^+^ memory T cells led to the isolation of additional clones from all patients tested, including P8 and P21. Collectively, these findings demonstrate that, in spite of its relatively small size, the RBD is recognized by CD4^+^ T cells of all individuals tested and accounts on average for 20% of the CD4^+^ T cell response to the Spike protein (**Fig. 2B**).

**Figure 2.**
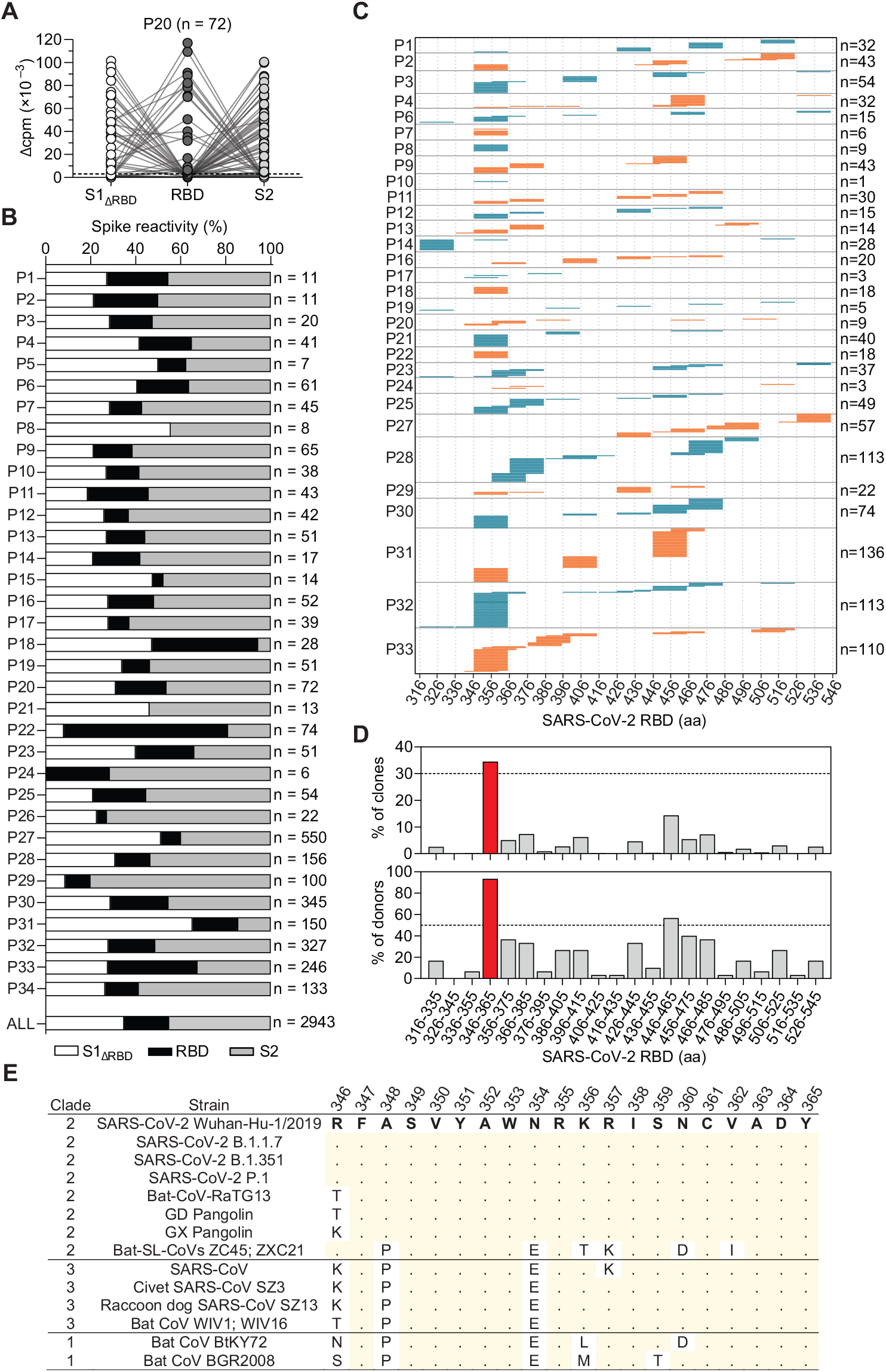
CD4^+^ T cell clones target multiple sites on the Spike protein. (**A, B**) CD4^+^ T cell clones (n = 2,943) were isolated from Spike-reactive cultures of 34 COVID-19 recovered patients and their specificity was mapped by stimulation with autologous B cells and 3 pools of peptides spanning S1ΔRBD, RBD, and S2, using as readout ^3^H-thymidine incorporation. (**A**) Characterization of representative T cell clones (n = 72) from patient P20. Proliferation was assessed on day 3 after a 16-h pulse with ^3^H-thymidine and expressed as counts per minute after subtraction of the unstimulated control value (Δcpm). (**B**) Percentage of T cell clones specific for S1ΔRBD (white), RBD (black), and S2 (grey) in the 34 patients tested; the number of clones tested is indicated for each patient. The distribution of all Spike-reactive T cell clones isolated from 34 COVID-19 patients (ALL, n = 2,943) is also indicated. (**C**) RBD-specific T cell clones (n = 1,149) isolated from 30 patients were further characterized for their epitope specificity using 15mer peptides overlapping by 10 spanning the S316-545 RBD sequence. The 20mer specificity of each clone is represented by a horizontal line and the total number of clones mapped for each patient is indicated on the right. (**D**) Percentage of clones specific and percentage of donors carrying T cells specific for different 20mer segments of the RBD. Data for the immunodominant region S346-365 is indicated in red. (**E**) Multiple sequence alignment of the immunodominant region S346-365 of SARS-CoV-2 with homologous sequences in different Sarbecoviruses, Human and animal SARS-related coronaviruses are arranged by clades. Amino acid numbers of SARS-CoV-2 Spike protein are reported on top. Amino acid substitutions compared to SARS-CoV-2 Wuhan-Hu-1/2019 reference strain are reported; dots indicate identity to SARS-CoV-2 reference strain.

Using a matrix-based approach, we mapped the epitope specificity of 1,149 RBD-reactive CD4^+^ T cell clones isolated from 30 patients (**Fig. 2C**). This analysis showed that in each patient T cell clones recognize multiple sites that altogether covered almost all the RBD sequence. Individual patterns were clearly evident, most likely reflecting restriction by different HLA class II molecules. However, certain regions emerged as immunodominant, such as those spanning residues S346-385 and S446-485 (**Fig. 2C**). Strikingly, a 20 amino acid region (S346-365) was recognized by 93% of the donors (28 out of 30) and by 34% of the clones (396 out of 1,149) (**Fig. 2D**). This region is highly conserved among human Sarbecoviruses, including the recently emerged variants of concern (VOC), and zoonotic Sarbecoviruses (**Fig. 2E**) (*18*). RBD-specific and S346-365-specific clones were found in the Tcm, Tem and cTfh subsets, consistent with priming of effector and memory T cells (**Fig. S3A, B**). Collectively, these findings demonstrate that RBD is highly immunogenic in vivo and identify a large number of naturally processed T cell epitopes including a conserved immunodominant region.

To dissect the CD4^+^ T cell response to the immunodominant S346-365 region, we sequenced TCR Vβ chains of 323 specific T cell clones. The 200 clonotypes identified used a broad spectrum of TCR Vβ genes and, even in the same donor, carried different CDR3 sequences (**Fig. 3A, B** and **Table S2**). The TCR Vβ sequences of T cell clones specific for the S346-365 region isolated from patients P31 and P33 were also detected in the repertoire of total CD4^+^ memory T cells immediately after isolation from peripheral blood and some of them were among the top 5% clonotypes (**Fig. 3C**).

**Figure 3.**
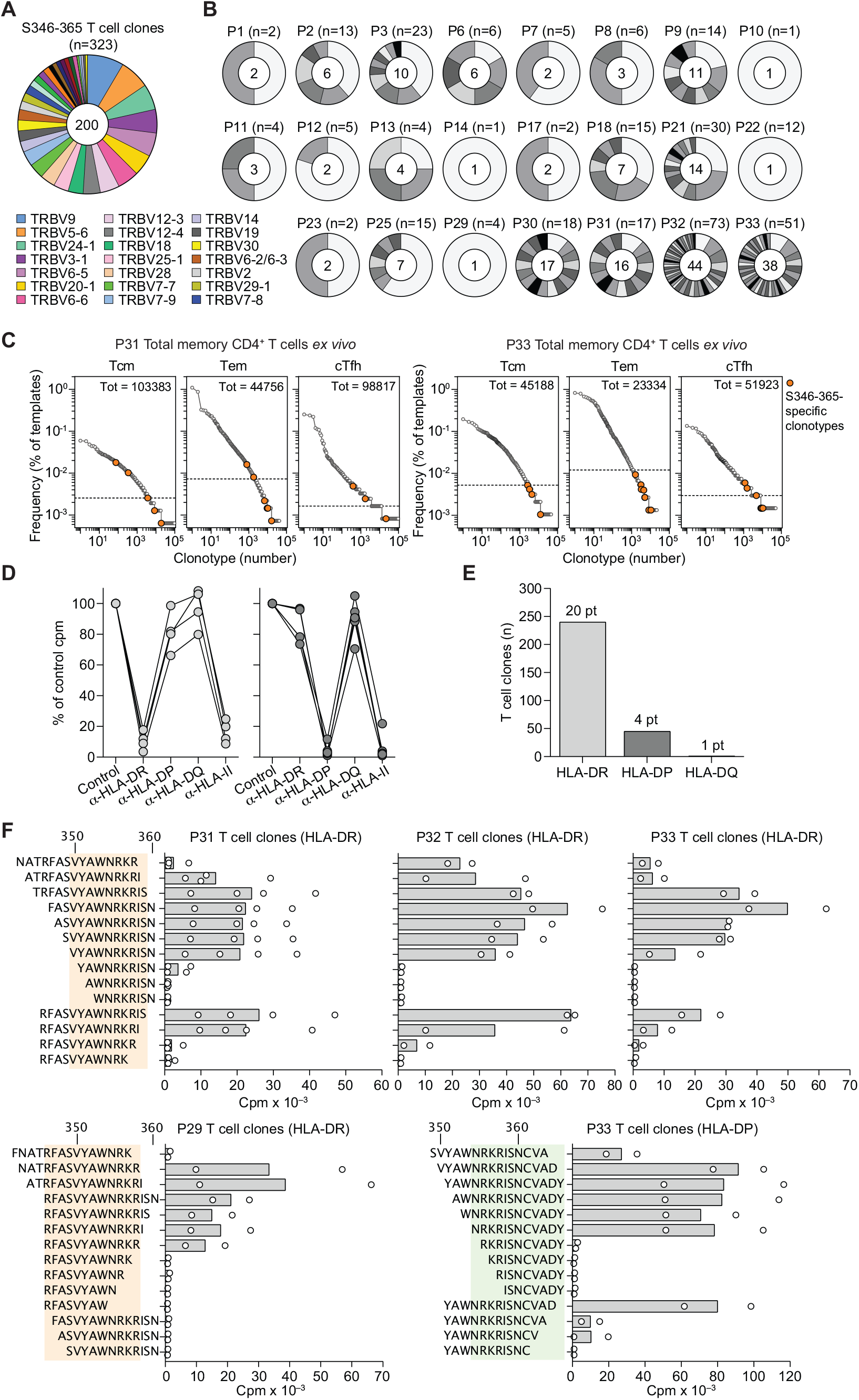
The immunodominant S346-365 RBD region contains nested peptides targeted by T cells restricted by HLA-DR and HLA-DP. (**A, B**) Rearranged TCR Vβ sequences of S346-365-reactive CD4^+^ T cell clones (n = 323) isolated from 23 patients were determined by RT-PCR followed by Sanger sequencing. (**A**) TCR Vβ gene usage in the 200 unique clonotypes identified from S346-365-reactive T cell clones. Each slice of the chart indicates a different Vβ gene. The size of each slice is proportional to the number of unique TCR Vβ clonotypes using that particular Vβ gene. Color-coded legend is reported for the top 21 Vβ genes (used by at least 4 different TCR Vβ clonotypes). (**B**) Number of S346-365-reactive T cell clones and clonotypes identified in the 23 patients. Each slice of the chart indicates a different TCR Vβ clonotype. The size of each slice is proportional to the number of sister clones bearing the same TCR Vβ sequence. The number of clones analyzed from each patient is reported on top of each pie chart; the number of TCR Vβ clonotypes identified is reported at the center. (**C**) Frequency distribution of TCR Vβ clonotypes from CD4^+^ Tcm Tem and cTfh subsets sequenced directly after ex vivo isolation from patients P31 and P33. Colored circles mark the TCR Vβ clonotypes found among the S346-365-specific T cell clones isolated from the same patient. Dotted lines in the graphs indicate the frequency threshold of the top 5% expanded clonotypes. (**D**) HLA class II isotype restriction of S346-365-specific T cell clones (n = 10) isolated from P33 was determined by stimulation with peptide-pulsed autologous APCs in the absence (control) or in the presence of blocking antibodies to HLA-DR, -DP, -DQ or pan HLA class II. Proliferation was assessed on day 3 after a 16-h pulse with ^3^H-thymidine. Data are expressed as percentage of control counts per minute (Cpm). (**E**) Distribution of HLA class II isotype restrictions of S346-365-reactive CD4^+^ T cell clones (n = 286) isolated from 22 patients, as determined by >80% inhibition of proliferation. (**F**) The minimal peptide recognized by S346-365-reactive CD4^+^ T cell clones (n = 12) isolated from 4 patients was mapped by stimulation with autologous APCs pulsed with a panel of truncated peptides. Proliferation was assessed on day 3 after a 16-h pulse with ^3^H-thymidine and expressed as counts per minute (Cpm). The minimal amino acid sequences recognized by T cell clones are highlighted with colored shading.

To determine the HLA class II isotype used by S346-365-specific T cell clones, we stimulated the clones in the presence of blocking antibodies to HLA-DR, -DP and -DQ (**Fig. 3D, E**). Most of the T cell clones analyzed (n = 240 from 20 patients) were HLA-DR-restricted, while the remaining (45 from 4 patients) were HLA-DP-restricted and one was HLA-DQ-restricted. The minimal peptide epitope recognized by HLA-DR- and HLA-DP-restricted T cell clones was mapped using a panel of truncated peptides (**Fig. 3F**). Interestingly, while HLA-DR-restricted clones from P31, P32 and P33 recognized the same 10 amino acid peptide (VYAWNRKRIS), HLA-DR-restricted clones from P29 recognized a different 12 amino acid peptide (RFASVYAWNRKR) and HLA-DP-restricted clones from P33 recognized an 11 amino acid peptide (NRKRISNCVAD). Collectively, these findings indicate that the S346-365 region comprises at least three nested epitopes recognized in association with different allelic forms of HLA-DR or HLA-DP by T cell clones that use a large set of TCR Vβ genes and CDR3 of different sequence and length.

To address the extent of T cell cross-reactivity between different Spike proteins, SARS-CoV-2 Spike-specific T cell lines from patients P28 and P33 were re-labelled with CFSE and stimulated with autologous monocytes pulsed with Spike proteins from other beta (SARS-CoV, HKU1 and OC43) or alpha (NL63 and 229E) coronaviruses. In these patients, a robust proliferation of SARS-CoV-2-specific T cell lines in the secondary cultures was observed in response to SARS-CoV and HKU1 (**Fig. 4A**). TCR Vβ sequencing of primary and secondary cultures identified a subset of cross-reactive clonotypes (**Fig. S4**). Out of 1,674 and 568 clonotypes present in the primary SARS-CoV-2-specific T cell lines from P28 and P33, 17% and 25% were found in SARS-CoV secondary cultures and 11% and 15% were found in HKU1 secondary cultures. Remarkably, 7% and 8% of the clonotypes were found in both SARS-CoV and HKU1 stimulated cultures suggesting the existence of T cell clones with broad specificity. To corroborate this finding, we isolated from secondary cultures several T cell clones that proliferated in response to 2 or even 3 different Spike proteins (**Fig. 4B** and **Table S3**). Importantly, the dose response curves indicate recognition of naturally processed peptides with similar functional avidity.

**Figure 4.**
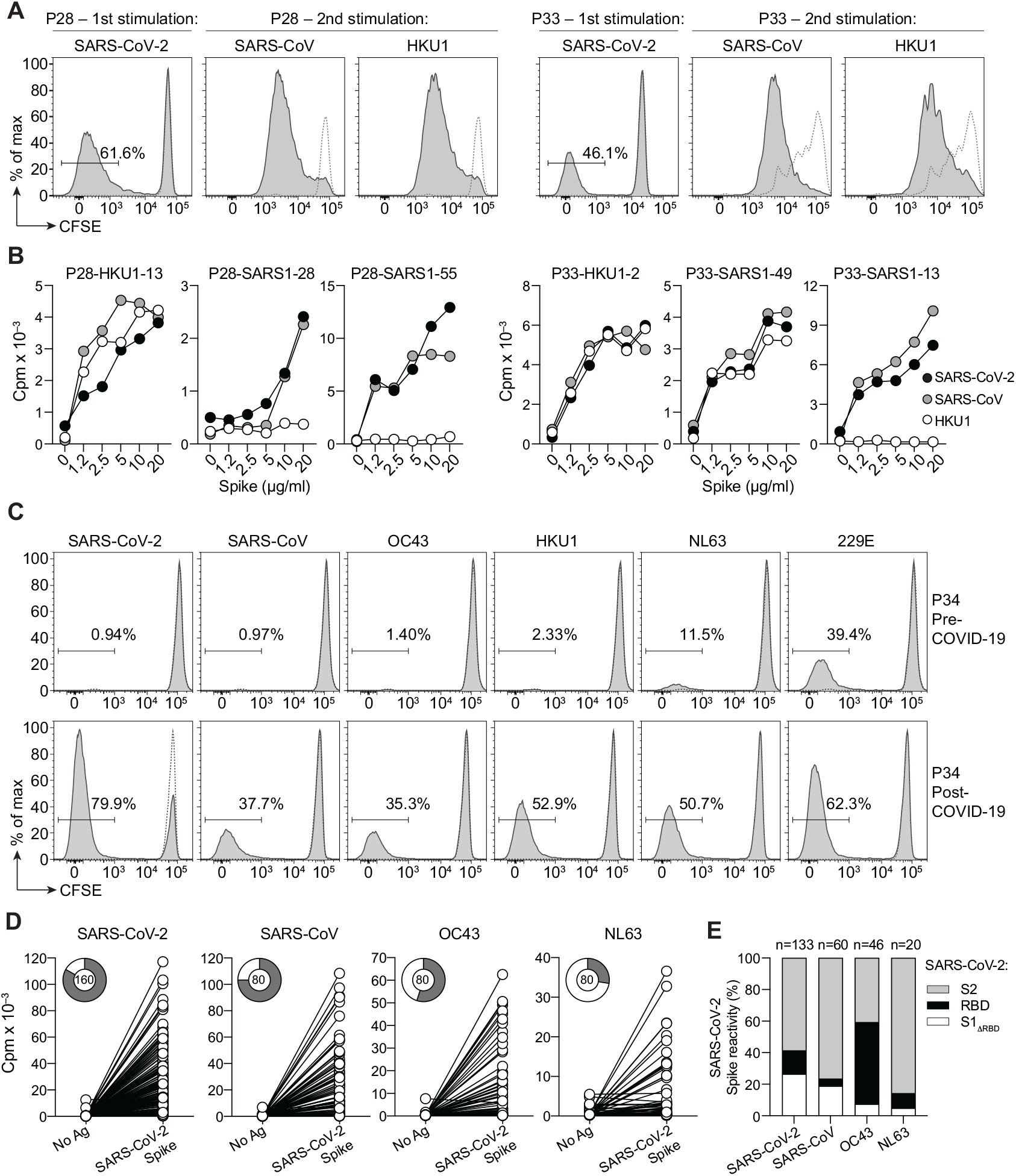
Identification of cross-reactive T cell clones by heterologous stimulations with Spike antigens. (**A**) CFSE-labelled CD4^+^ memory T cells were stimulated with recombinant SARS-CoV-2 Spike protein in the presence of autologous monocytes. CFSE^lo^ cells were expanded with IL-2 for 10 days, re-labelled and re-stimulated with SARS-CoV or HCoV-HKU1 Spike protein. Negative controls of T cells cultured with monocytes in the absence of antigen are reported as dashed lines. Shown are experiments performed with samples from patient P28 and P33. (**B**) Proliferative response of representative T cell clones isolated from P28 and P33 to autologous APCs pulsed with titrated doses of recombinant Spike proteins from SARS-CoV-2 (black), SARS-CoV (grey) or HCoV-HKU1 (white). (C) Proliferative response of memory CD4^+^ T cells isolated from P34 two years before and 1.5 months after COVID-19. CFSE-labelled cells were stimulated as above with different recombinant Spike proteins. (**D**) Proliferative response of patient P34 T cell clones isolated from the post-COVID-19 CFSE^lo^ fractions shown above. The T cell clones were stimulated with autologous APCs pulsed with a pool of peptides spanning the entire SARS-CoV-2 Spike protein. Pie charts show the total number of clones tested and the fraction of responsive clones. (**E**) Reactivity of T cell clones isolated from each culture (in **D**) was mapped by stimulation with autologous APCs pulsed with pools of peptides spanning the S1ΔRBD (white), RBD (black), and S2 (grey) regions of SARS-CoV-2 Spike. The histograms show the percentage of clones specific for each region. Total number of clones tested is indicated on top.

To ask whether cross-reactive T cells could be detected in pre-COVID-19 samples and could be expanded early on following SARS-CoV-2 infection, we analyzed the T cell response in a COVID-19 recovered patient from whom we had previously cryopreserved PBMCs. Memory CD4^+^ T cells from pre- and post-COVID-19 samples were stimulated with a panel of recombinant Spike proteins (**Fig. 4C**). In this individual, a robust T cell proliferation in the pre-COVID-19 sample was detected primarily against alpha coronaviruses NL63 and 229E, while the response to beta coronaviruses HKU1 and OC43 was limited and the response to SARS-CoV and SARS-CoV-2 was undetectable. Conversely, in the post-COVID-19 sample a strong T cell proliferation was observed not only in response to SARS-CoV-2, but also in response to all other alpha and beta coronavirus Spike proteins. This was accompanied by the appearance of antibodies against SARS-CoV-2 as well as an increase in serum antibody titers against HKU1 and OC43 (J.S.L., unpublished data).

The striking increase in the response to unrelated Spike proteins in the post-COVID-19 sample may be caused by the recall of pre-existing cross-reactive memory T cells or the priming of cross-reactive T cells. To measure the extent of cross-reactivity, we isolated T cell clones from SARS-CoV-2-, SARS-CoV-, OC43- or NL63-reactive memory T cell lines and found that a variable proportion of these clones proliferated in response to a SARS-CoV-2 Spike peptide pool (**Fig. 4D**). The specificity of cross-reactive T cell clones was further mapped using peptide pools to the different SARS-CoV-2 Spike domains and was found to be primarily directed against the S2 domain but to comprise also T cell clones reactive to the S1 or RBD domain (**Fig. 4E**). Collectively, these findings identify, at the clonal level, an extensive CD4^+^ T cell cross-reactivity among coronavirus Spike proteins, which primarily targets the S2 domain, consistent with its higher degree of sequence identity (*19-21*).

Using naturally processed antigens for the isolation of almost 3,000 CD4^+^ T cell clones, we unbiasedly mapped the T cell epitopes recognized by patients recovered from COVID-19. At variance with initial reports, we document a robust CD4^+^ T cell response to the RBD and identified a conserved immunodominant region encompassing residues S346-365 in the RBD that is recognized by virtually all donors, a finding that is relevant for vaccine design since the RBD is also the target of most neutralizing antibodies (*16, 17*). These findings were not anticipated in previous studies based on bioinformatics predictions (*2, 3*) and short-term peptide stimulation of bulk PBMCs, highlighting the value and suitability of classical unbiased T cell stimulation and cloning approach for the dissection of antigen-specific T cell repertoires.

The reason for the immunodominance of RBD S346-365 may be due to the presence of at least three nested T cell epitopes presented by HLA-DR and HLA-DP as shown in this study, but may also be due to the relative abundance of naturally processed peptides, as previously observed in the context of the immunodominant response to influenza hemagglutinin (*13*). Indeed, peptides spanning the immunodominant regions here identified were also detected by HLA class II immunopeptidomics in dendritic cells pulsed with SARS-CoV-2 Spike protein (*22*). Interestingly, the immunodominant S346-365 region is conserved among different Sarbecoviruses and has not been found mutated in the new SARS-CoV-2 VOC isolates. Furthermore, this region is also a contact site for a broadly reactive neutralizing antibody (*23*), providing a striking example of convergence of T and B cells around a conserved epitope.

Sequencing of TCR Vβ locus in T cell clones and polyclonal cultures reveals that SARS-CoV-2 infection primes a large CD4^+^ T cell response comprising inflammatory and helper T cells, as well as precursors of long-lived memory T cells. In this primary response, it was interesting to find evidence for an extensive clonal diversification towards different fates and functions, as previously demonstrated in mice and in in vitro primed human naïve T cells (*14, 15*). Furthermore, the detection of a large repertoire of SARS-CoV-2 reactive cTfh cells is indicative of robust germinal center reaction, which is necessary for the deployment of high affinity antibodies (*24*). Follow-up studies on clonotype dynamics and persistence will allow to dissect the development of T cell memory to a relevant pathogen and to test different models of memory generation (*25*).

Our results also highlight the extensive cross-reactivity of Spike-specific T cells that can be documented in vivo and not only between the closely related Sarbecoviruses (SARS-CoV-2 and SARS-CoV) but also with the more divergent endemic coronaviruses. The availability of a large number of cross-reactive T cell clones is not only instrumental to define target sites in relevant pathogens but also to understand whether cross-reactivity is due to epitope structural similarities or to TCR binding degeneracy (*11, 26*).

While the narrow specificity of neutralizing antibodies can readily select escape mutants, the extensive sequence coverage and breadth observed in the CD4^+^ T cell response to SARS-CoV-2 Spike antigen offers an opportunity for vaccine design. The possibility of leveraging a robust cross-reactive T helper cell function against conserved sites will be instrumental to drive neutralizing antibody responses to adaptive vaccines that incorporate escape mutations found in emerging SARS-CoV-2 variants.

## Materials and Methods

### Study subjects

Blood samples were obtained from a cohort of COVID-19 convalescent patients collected between June 8, 2020 and November 24, 2020 and from healthy donors collected and cryopreserved in summer 2019 (obtained from the Swiss Blood Donation Center of Basel and Lugano, pre-pandemic samples) or on November 2018 (P34). The study protocol was approved by the Cantonal Ethics Committee of Ticino, Switzerland (CE-TI-3428, 2018-02166). All blood donors provided written informed consent for participation in the study. Human primary cell protocols were approved by the Federal Office of Public Health (no. A000197/2 to F.S.). Illness of SAoV-2 infected individuals was categorized according to the CDC guidelines (https://www.covid19treatmentguidelines.nih.gov/overview/clinical-spectrum/) as: i) asymptomatic if they tested positive for SARS-CoV-2 and reported any COVID-19-related symptoms; ii) mild if they reported any of the following symptoms: fever, cough, sore throat, malaise, headache, muscle pain, nausea, vomiting, diarrhea, loss of taste and smell, but no shortness of breath, dyspnea, or abnormal chest imaging; iii) moderate if evidence of lower respiratory disease during clinical assessment or imaging and with oxygen saturation (SpO2) ≥94% on room air; iv) severe if SpO2 <94% on room air.

### Cell purification and sorting

Peripheral blood mononuclear cells (PBMCs) were isolated by Ficoll-Paque Plus (GE Healthcare) gradient. CD14^+^ monocytes and total CD4^+^ T cells were isolated by positive selection using CD14 and CD4 magnetic microbeads, respectively (Miltenyi Biotech). Total CD4^+^ cells were stained at 37°C for 15 min with a primary anti-human-CXCR5 (clone 51505; cat. no. MAB190) from Bio-Techne, followed by staining with a biotinylated secondary goat anti-mouse IgG2b (cat. no. 1090-08) from Southern Biotech. After washing, cells were stained with PE/Cy7-Streptavidin (cat. no. 405206) from BioLegend, and with the following fluorochrome-labeled mouse monoclonal antibodies: CD8-PE-Cy5 (clone B9.11; cat. no. A07758), CD56-PE/Cy5 (clone N901; cat. no. A07789) from Beckman Coulter, CD25-PE (clone M-A251; cat. no. 555432) from BD Biosciences, CD4-PE–Texas Red (clone S3.5; cat. no. MHCD0417), CD45RA-Qdot 655 (clone MEM-56; cat. no. Q10069) from ThermoFisher Scientific, CCR7-BV421 (clone G043H7; cat. no. 353208) from BioLegend. Memory CD4^+^ T cells were sorted to over 98% purity on a FACSAria III (BD) after exclusion of CD8^+^, CD56^+^, CD25^br^ cells and CD45RA^+^CCR7^+^) naïve CD4^+^ T cells. In some experiments, memory CD4^+^ T cells were divided in cTfh (sorted as CXCR5^+^ cells), Tcm (sorted as CCR7^+^CXCR5^–^ cells) and Tem (sorted as CCR7^–^CXCR5^–^ cells).

### Antigens and peptides

The following recombinant proteins were purchased from Sino Biological Inc: SARS-CoV-2 (2019-nCoV) Spike protein (S1+S2 ECD, cat.no. 40589-V08B1), SARS-CoV-2 (2019-nCoV) Nucleocapsid protein (cat.no. 40588-V08B), SARS-CoV (S577A, Isolate Tor2) Spike protein (S1+S2 ECD, cat.no. 40634-V08B), HCoV-HKU1 (isolate N5) Spike protein (S1+S2 ECD, cat.no. 40606-V08B). Peptides were synthesized as crude material on a small scale (1 mg) by Pepscan (Lelystad, The Netherlands). Peptides used in the study included 15mers overlapping of 10 covering the entire sequences of SARS-CoV-2 Spike protein (UniProtKB: P0DTC2; 253 peptides).

### Cell culture and T cell stimulation

T cells were cultured in RPMI 1640 medium supplemented with 2 mM glutamine, 1% (vol/vol) nonessential amino acids, 1% (vol/vol) sodium pyruvate, penicillin (50 U/ml), streptomycin (50 µg/ml) (all from Invitrogen) and 5% human serum (Swiss Red Cross). Sorted memory CD4^+^ T cells were labelled with 5-(and 6)-carboxyfluorescein diacetate succinimidyl ester (CFSE, ThermoFisher) and cultured at a ratio of 2:1 with irradiated autologous monocytes untreated or pulsed for 3 h with recombinant SARS-CoV-2 Spike protein (2.5 μg/ml). After 6 days, cells were stained with antibodies to CD25-PE (clone M-A251; cat. no. 555432) from BD Biosciences and ICOS-APC (clone C398.4A; cat. no. 313510) from BioLegend. Cytokine concentrations in the day 6 supernatants of antigen-stimulated cultures were assessed by Luminex bead-based assay (Thermo Fisher Scientific) according to the manufacturer’s instructions. Proliferating, activated T cells were FACS-sorted as CFSE^lo^CD25^+^ICOS^+^, expanded in vitro in the presence of IL-2 (500 IU/ml) and cloned by limiting dilution. In some experiments, CFSE^lo^ cultures were relabeled with CFSE and re-stimulated with antigen-pulsed irradiated autologous monocytes; readout of T cell proliferation was determined at day 4-5 after secondary stimulation.

### Isolation and characterization of T cell clones

T cell clones were isolated from CFSE-low cells by limiting dilution using 1 μg/mL PHA-P, 1000U/ml IL-2 and allogeneic irradiated PBMCs. T cell clone reactivity was determined by stimulation with a set of 253 synthetic peptides (15mers overlapping of 10) spanning SARS-CoV-2 Spike protein (1.5 μM per peptide) in the presence of irradiated autologous monocytes or EBV-immortalized B cells as APCs. To determine the SARS-CoV-2 Spike domains targeted by T cell clones, the set of 253 peptides was grouped in 3 different pools spanning the following amino acid ranges: S1-325 and S536-685 (pool S1ΔRBD, comprising 91 peptides); S316-545 (pool RBD, comprising 44 peptides); S676-1273 (pool S2, comprising 118 peptides). To determine the minimal sequence targeted by SARS-CoV-2 RBD-reactive T cell clones, the 44 peptides comprising the pool RBD were subdivided into 5 horizontal and 5 vertical peptide pools to construct a RBD epitope mapping matrix. Epitope mapping of SARS-CoV-2 RBD was performed by stimulation of T cell clones with irradiated autologous APCs, untreated or pre-pulsed with the RBD epitope mapping matrix (1.5 μM per peptide). Specificity to a region of 20 amino acids was assigned upon reactivity to one of the 5 horizontal and one of the 5 vertical pools. In some experiments, SARS-CoV-2 Spike peptides were titrated by serial dilution. To determine HLA restriction, T cell clones were stimulated with autologous APCs pulsed with SARS-CoV-2 Spike peptides, in the absence or presence of blocking anti-class II monoclonal antibodies produced in house from hybridoma cell lines (anti-HLA-DR, clone L243 from ATCC, cat. no. HB-55; anti-HLA-DQ, clone SPVL3 (27); anti-HLA-DP, clone B7/21 (28), anti-pan-MHC-class-II, clone IVA12 from ATCC, cat. no. HB-145). In all experiments proliferation was assessed on day 3, after incubation for 16 h with 1 µCi/ml [methyl-^3^H]-thymidine (Perkin Elmer). Data were expressed as counts per min (Cpm) or as Δcpm after subtraction of control cultures performed in the absence of antigen.

### Sequence analysis of TCR Vβ genes

Sequence analysis of rearranged TCR Vβ genes of T cell clones was performed as previously described. Briefly, cDNA from individual T cell clones was obtained by reverse transcription of total RNA from 10^3^-10^4^ cells per reaction. Rearranged TCR Vβ genes were PCR amplified using forward primer pool targeting Vβ genes, and reverse primer pairing to C1–C2 β-chain constant region. Sequence amplification was assessed through agarose gel electrophoresis; successfully amplified fragments were sequenced by Sanger method, and TCR sequence annotation was carried out by using IMGT/V-QUEST algorithm (*29*).

### TCR Vβ deep sequencing

Ex vivo-sorted memory CD4^+^ T cell subsets and CFSE^lo^ fractions of antigen-stimulated memory CD4^+^ T cell cultures were analyzed by deep sequencing. In brief, 5×10^5^ T cells were centrifuged and washed in PBS and genomic DNA was extracted from the pellet using QIAamp DNA Micro Kit (Qiagen), according to manufacturer’s instructions. Genomic DNA quantity and purity were assessed through spectrophotometric analysis. Sequencing of TCR Vβ CDR3 was performed by Adaptive Biotechnologies using the ImmunoSEQ assay (http://www.immunoseq.com). In brief, following multiplex PCR reaction designed to target any CDR3 Vβ fragments; amplicons were sequenced using the Illumina HiSeq platform. Raw data consisting of all retrieved sequences of 87 nucleotides or corresponding amino acid sequences and containing the CDR3 region were exported and further processed. The assay was performed at deep level for ex vivo-sorted memory CD4^+^ T cell subsets (detection sensitivity, 1 cell in 200,000) and at survey level for CFSE^lo^ antigen-reactive cultures (detection sensitivity, 1 cell in 40,000). Each TCR Vβ clonotype was defined as the unique combination of a productively rearranged CDR3 amino acid sequence and its related V and J genes (bioidentity); data processing was done using the productive frequency of templates provided by ImmunoSEQ Analyzer V.3.0 (http://www.immunoseq.com). For each repertoire, a frequency corresponding to the top 5th percentile in the frequency-ranked list of unique clonotypes was chosen as threshold (top 5%). Chao-Jaccard overlap between pairs of TCR Vβ repertoires was calculated using R package “fossil” (*30*).

### Conservation analysis

Accession numbers of Spike protein sequences used for S346-365 conservation analysis are: SARS-CoV-2 Wuhan-Hu-1/2019 (GenBank: NC_045512.2); SARS-CoV-2 B.1.1.7 (GISAID: EPI_ISL_700654); SARS-CoV-2 B.1.351 (GISAID: EPI_ISL_700492); SARS-CoV-2 P.1 (GISAID: EPI_ISL_833137); Bat-CoV-RaTG13 (GenBank: QHR63300.2); GD Pangolin (GISAID: EPI_ISL_410721; 471470; 471469; 471468; 471467); GX Pangolin (GISAID: EPI_ISL_410543; 410542; 410541; 410540; 410539); Bat-SL-CoV ZC45 (GenBank: MG772933.1); Bat-SL-CoV ZXC21 (GenBank: MG772934.1); SARS-CoV (GenBank: Urbani AAP13442.1; Tor2 NC_004718.3; TW1 AAP37015.1; P2 ACQ82725.1; Frankfurt1 AAP33696.1); Civet SARS-CoV SZ3 (GenBank: AY304486.1); Raccoon dog SARS-CoV SZ13 (GenBank: AY304487.1); Bat CoV WIV1 (GenBank: AGZ48828.1); Bat CoV WIV16 (GenBank: KT444582.1); Bat CoV BtKY72 (GenBank: KY352407.1); Bat CoV BGR2008 (GenBank: GU190215.1). Clades were defined according to Lu et al. (*18*).

### Statistical analysis

Statistical analyses were performed using GraphPad Prism 8 software. Significance was assigned at *P* value < 0.05, unless stated otherwise. Specific tests are indicated in the figure legends for each comparison. Analysis of TCR Vβ repertoires was performed using R software version 3.5.1.

## Acknowledgements

We thank all participants to the study and the personnel at the hospitals for blood collection, Greta Durini for cell isolation, Daniele Lilleri and the Tipizzazione Lab of the IRCCS San Matteo Hospital Foundation San Matteo Pavia, Italy, for HLA typing.

## Funding

The study was in part financed by the Henry Krenter Foundation, the Swiss National Science Foundation (grant n. 189331) and the EOC research funds. FS and the Institute for Research in Biomedicine are supported by the Helmut Horten Foundation.

## Author contributions

ACa and FS contributed to study concept and design. JSL, DV, FM contributed to experimental work. DJ performed cell sorting. SJ provided technical support. MF performed bioinformatics analysis. TT, AFP, MB, CG, PF, ACe contributed clinical samples. ACa, AL and FS analyzed the data and wrote the manuscript. All authors contributed to interpretation of data and critical revision of the manuscript.

## Competing interests

Authors declare no competing interests.

## Data and materials availability

TCR Vβ sequences have been deposited in the ImmunoAccess database and an open access link will be provided in due time. All other data is available in the main text or the supplementary materials.

**Fig. S1.**
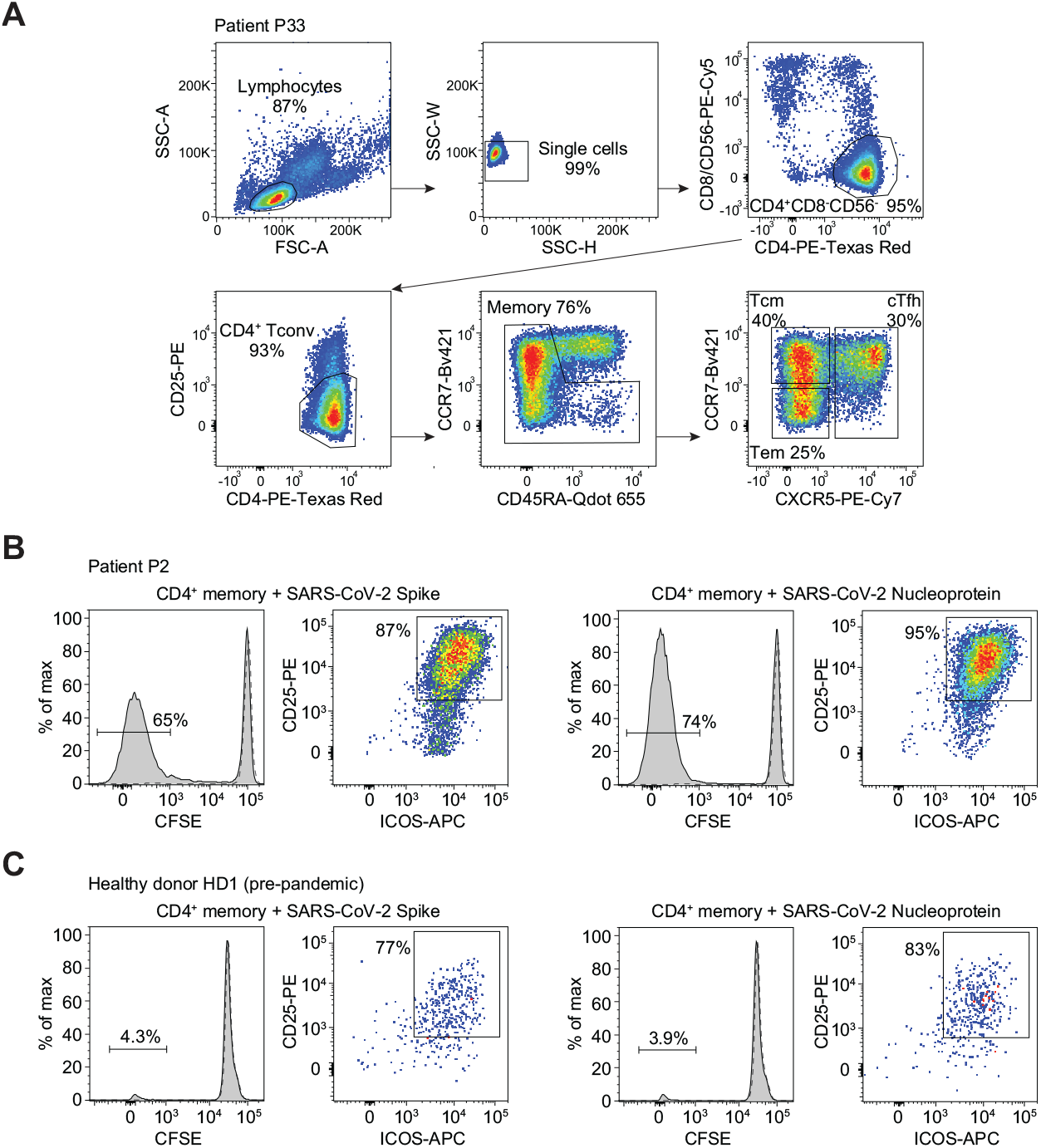
Sorting of T cell subsets and identification of SARS-CoV-2 Spike- and Nucleoprotein-reactive CD4^+^ T cells in COVID-19 and pre-pandemic samples. (**A**) Sorting strategy to isolate CD4^+^ total memory T cells and Tcm, Tem and cTfh subsets. (**B, C**) Characterization of antigen-specific T cells by CFSE dilution combined with CD25 and ICOS co-expression at day 7 following stimulation with Spike or Nucleoprotein in the presence of autologous monocytes. Negative controls of T cells cultured with monocytes alone are reported as dashed lines. Shown are data from patient P2 and from a pre-pandemic healthy donor sample (HD1).

**Fig. S2.**
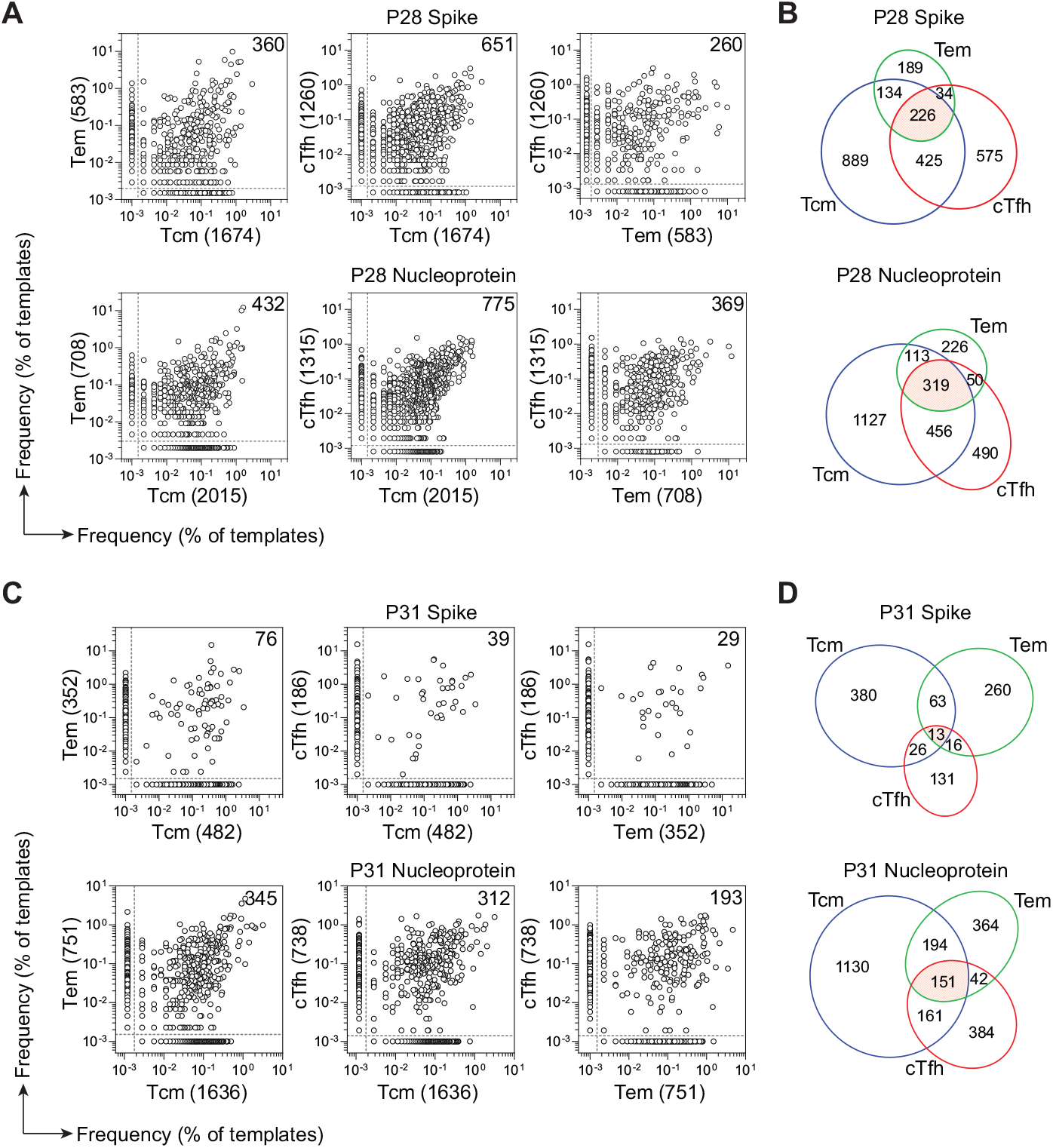
Repertoire analysis of SARS-CoV-2 Spike- and Nucleoprotein-specific T cells isolated from memory CD4^+^ T cell subsets. (**A, C**) Pairwise comparison of TCR Vβ clonotype frequency distribution of Spike- or Nucleoprotein-specific Tcm, Tem and cTfh cells from patient P28 (**A**) and P31 (**C**). Frequencies are shown as percentage of productive templates. Total number of clonotypes is indicated in the *x* and *y* axis. Values in the upper right corner represent the number of clonotypes shared between two samples. (**B, D**) Venn diagram showing the number of unique and shared Spike- or Nucleoprotein-reactive clonotypes between Tcm, Tem and cTfh subsets of patient P28 (**B**) and P31 (**D**). The red area represents clonotypes identified in all 3 subsets.

**Fig. S3.**
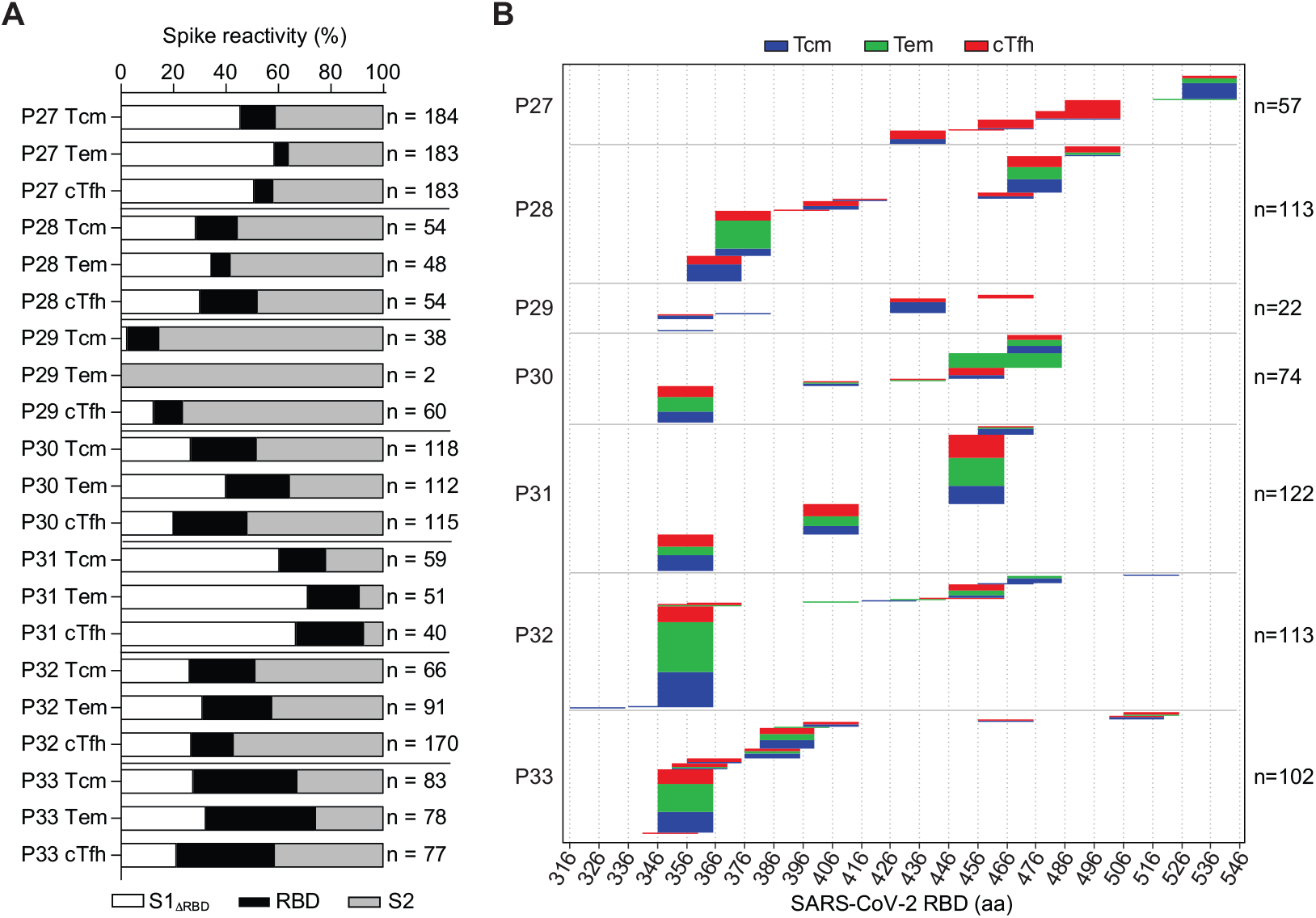
Epitope mapping of SARS-CoV-2 Spike specific T cell clones isolated from memory T cell subsets. (**A**) T cell clones were isolated from Spike-reactive cultures of Tcm, Tem and cTfh cells as in Fig. 2 and their specificity was mapped using autologous APCs pulsed with pools of peptides spanning the S1ΔRBD (white), RBD (black), and S2 (grey) regions of SARS-CoV-2 Spike. (**B**) RBD-specific T cell clones were further stimulated using a matrix of 15mer peptides overlapping of 10. Each line represents the specificity of a clone mapped to a 20 amino acid region. The colors indicate the subset of origin of each clone; the number of clones analyzed for each patient is indicated on the right.

**Fig. S4.**
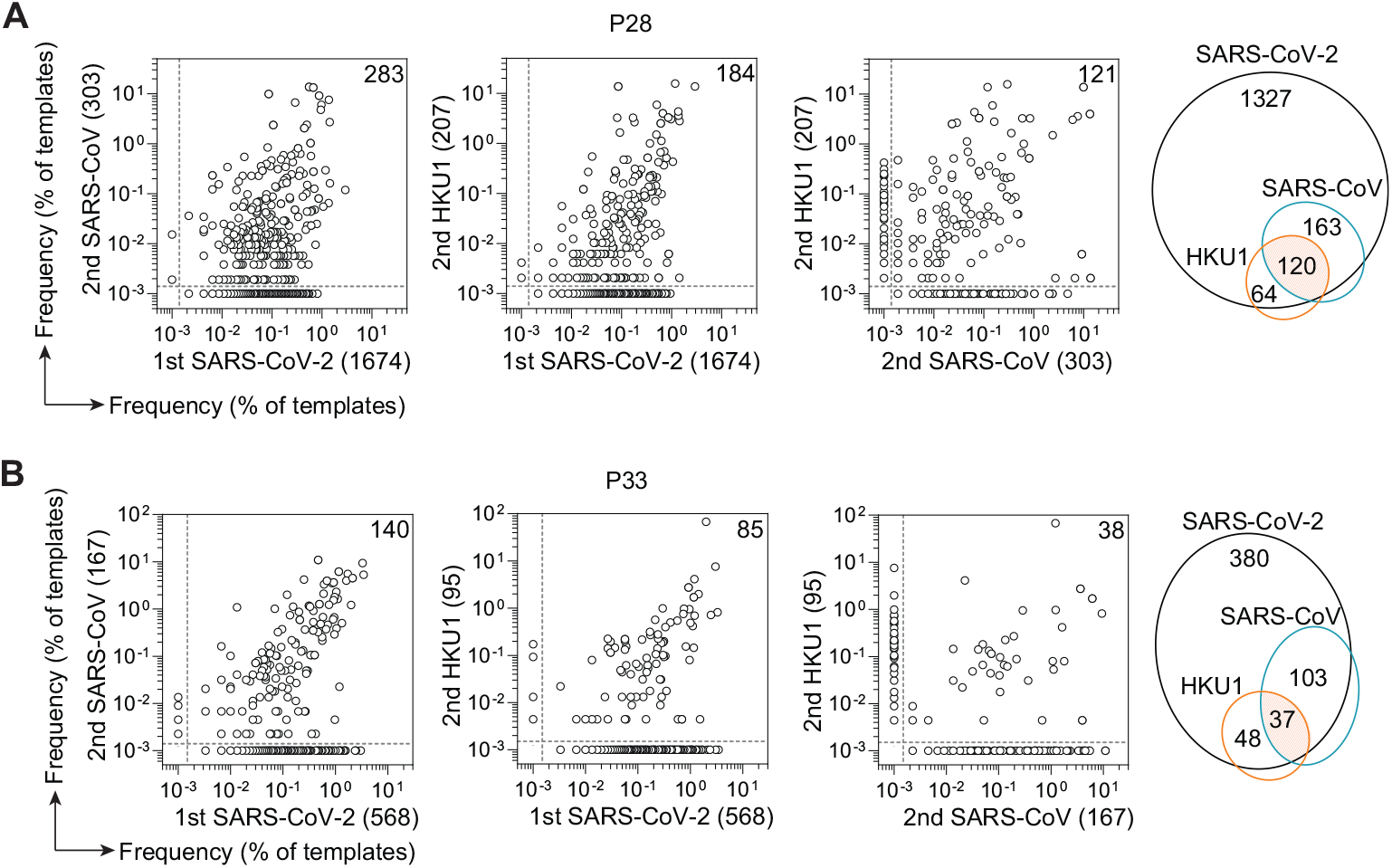
Repertoire analysis of cross-reactive CD4^+^ T cells. (**A, B**) Dot plots show for patient P28 (**A**) and P33 (**B**) the shared TCR Vβ clonotypes between sorted CFSE^lo^ T cells isolated following primary stimulation with SARS-CoV-2 Spike protein and sorted CFSE^lo^ T cells isolated following secondary re-stimulation with SARS-CoV or HCo-HKU1 Spike proteins. Frequencies are shown as percentage of productive templates. Total number of clonotypes is indicated in the *x* and *y* axis. Values in the upper right corner represent the number of clonotypes shared between two samples. The Venn diagrams show the number of clonotypes shared among different Spike-reactive cultures.

**Table S1.**
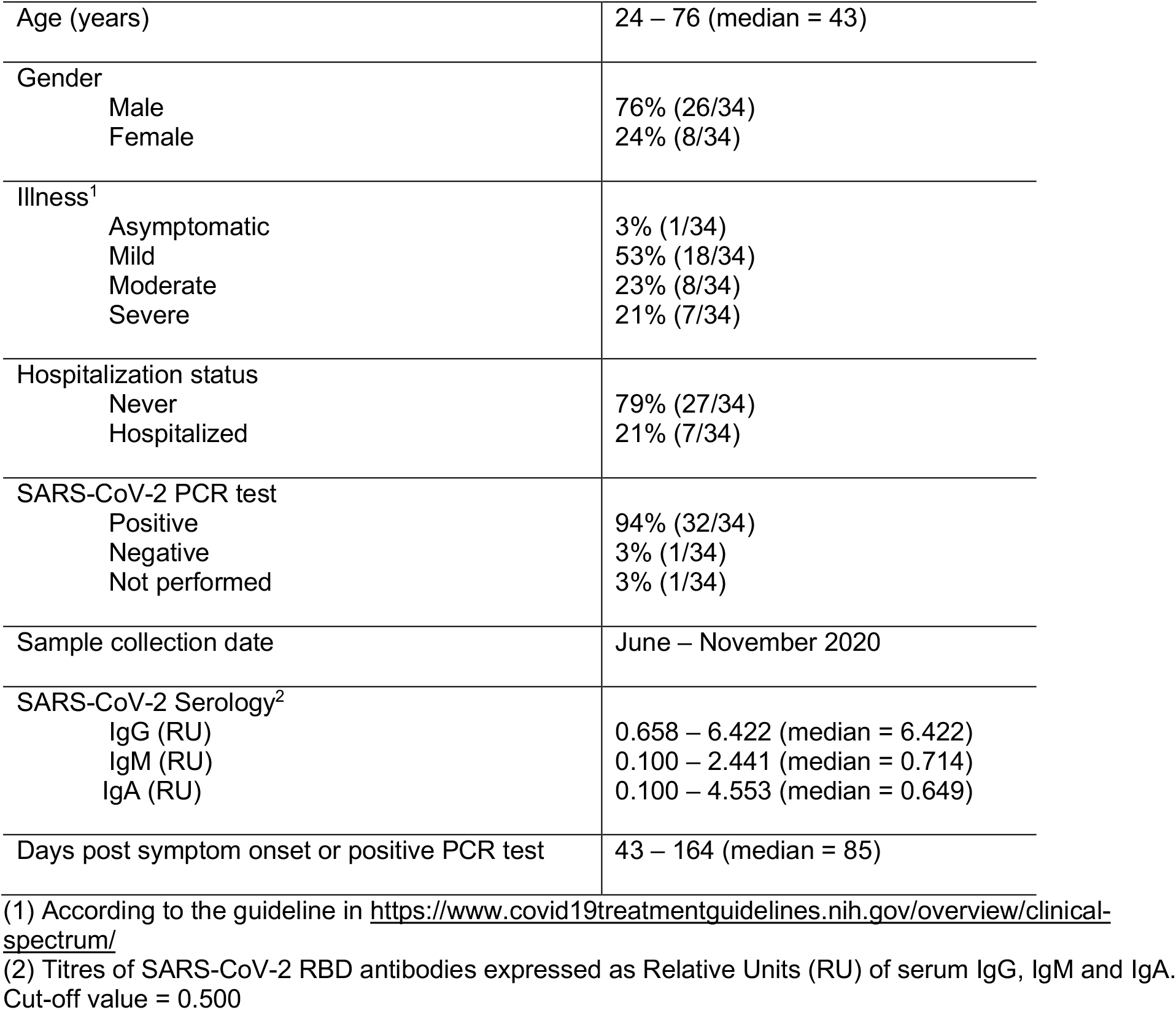
Characteristics of study participants (n = 34)

**Table S2.**
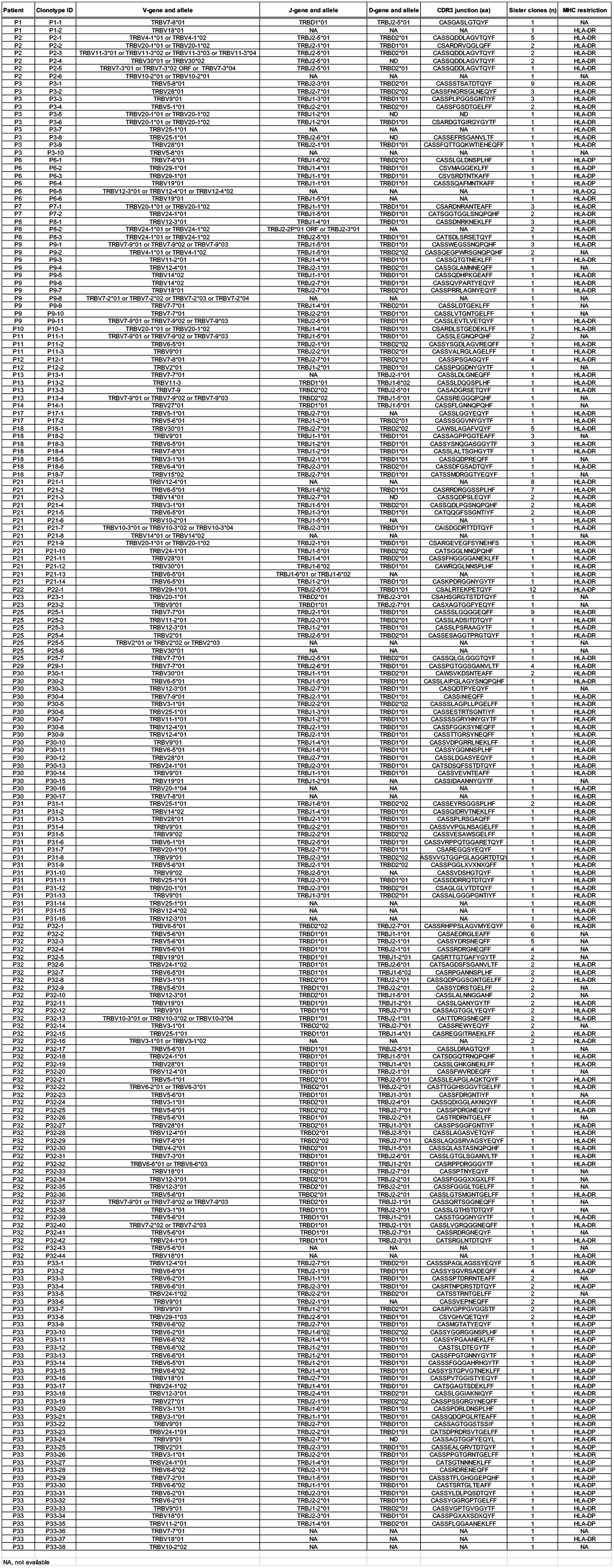
TCR Vβ sequence and MHC restriction of S346-365 specific T cell clones.

**Table S3.**
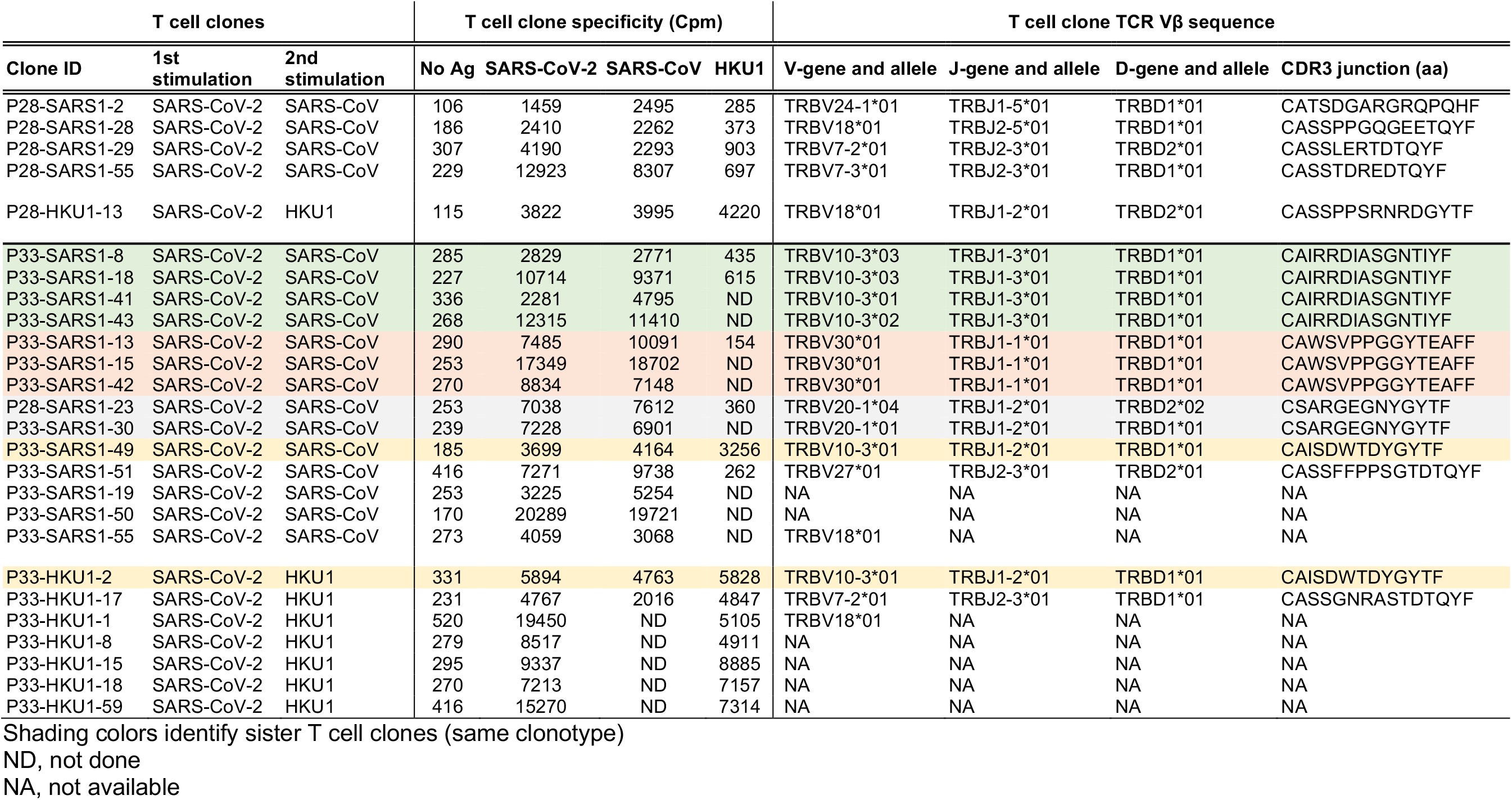
Specificity and TCR Vβ sequence of cross-reactive T cell clones.

## Notes

### Competing Interest Statement

The authors have declared no competing interest.

